# Silent Footprints of Ebolavirus in the Forest: Serological Clues from Wild Non-Human Primates in the Democratic Republic of Congo

**DOI:** 10.64898/2026.03.09.710450

**Authors:** Charles Kumakamba, Clément Labarrere, Inestin Amona, Joa Mangombi-Pambou, Jean-Jacques Muyembe-Tamfum, Florence Fenollar, Oleg Mediannikov

## Abstract

Filoviruses, particularly Ebola virus (EBOV), remain a major public health concern in Central Africa. However, their circulation in wildlife during inter-epidemic periods remains poorly documented. Non-human primates (NHPs) may serve as ecological sentinels of viral dynamics at human-forest interfaces, yet surveillance is constrained by the limitations of invasive sampling.

We conducted a non-invasive investigation of EBOV exposure in free-ranging NHPs from the Mabali Forest Reserve (Equateur Province, Democratic Republic of Congo) for Ebola virus disease. A total of 630 fecal samples were collected and screened for active infection by PCR targeting the EBOV nucleoprotein gene; all samples tested negative. Molecular identification of host species was achieved in 569 samples (90.3%).

Fecal serology using an automated capillary western blot platform (JESS), targeting EBOV nucleoprotein, glycoprotein and viral protein 40 antigens, identified four seropositive individuals (0.70%), including two *Cercopithecus ascanius* and two *C. wolfi*. The detection of discrete immunoreactive bands consistent with EBOV-specific antibodies suggests prior exposure despite the absence of active outbreaks.

These findings provide the first serological evidence compatible with EBOV exposure in these two *Cercopithecus* species and support the hypothesis of low-level or cryptic viral circulation in forest ecosystems. The study highlights the feasibility and value of fecal serology as a non-invasive One Health surveillance tool for monitoring zoonotic pathogens at wildlife–human interfaces.

## 1. Introduction

Filoviruses, particularly Ebola virus (EBOV), are responsible for severe hemorrhagic fevers with case fatality rates that can reach very high levels in humans (1). Since their discovery in 1976, these viruses have caused multiple outbreaks across Central and West Africa (2), with major impacts on public health systems, local economies, and ecosystems (3). These emergence events are closely linked to complex ecological dynamics at the human– animal–environment interface, where certain mammalian species, especially non-human primates (NHPs) and bats, play a pivotal role in the silent or cryptic circulation of filoviruses (4)(5).

Moreover, the natural cycle of filoviruses, their circulation dynamics within ecosystems, and the precise identification of their reservoirs remain insufficiently elucidated. Although several lines of evidence suggest an important role for bats (order Chiroptera) as potential hosts and reservoirs of Ebola virus, direct proof of a definitive reservoir remains limited, and the ecological mechanisms enabling viral maintenance in wildlife are still poorly understood. Complex interactions among wildlife species, including NHPs, may facilitate interspecies transmission events, playing a key role in viral amplification and in the risk of zoonotic spillover to human populations. A better understanding of these ecological processes and host networks therefore represents a major challenge for anticipating future emergences and strengthening integrated surveillance strategies.

Despite recent technological advances, detecting filoviruses in natural environments remains a major scientific challenge. Field investigations suggest that viral circulation is difficult to document in wildlife populations, owing to temporal fluctuations in viral signals (6)(7), the low probability of detecting viral RNA in excreta using molecular tools (8)(9), and the existence of sporadic yet genuine subclinical infections in both humans and wildlife (10)(11)(12). Molecular approaches alone are therefore insufficient to reveal past or unexpected exposure events, which underscores the importance of integrating serological tools as indispensable complementary methods within a One Health surveillance framework (6).

It is therefore essential to develop surveillance strategies capable of detecting not only active infections but also weak or residual immunological signals indicative of silent viral circulation within ecosystems. Although still rarely used for filovirus surveillance, fecal serology appears particularly well suited to field constraints, as it allows sampling of free-ranging individuals without capture, stress, or disruption of natural behavior.

Non-human primates represent a particularly relevant model in this context. They share close phylogenetic, behavioral, and eco-epidemiological characteristics with humans, potentially acting as reservoirs, amplifiers, or ecological sentinels of viral circulation (13)(14). However, most previous serological studies have been conducted on captive, deceased, or outbreak-related individuals, limiting the representativeness of the data obtained (15).

Invasive sampling methods in NHPs present major ethical, logistical, and behavioral constraints, making their large-scale application in forested environments difficult. In response to these limitations, fecal serology coupled with high-precision analytical techniques, such as automated western blotting, has emerged as a promising alternative, enabling the sampling of free-ranging, healthy, and undisturbed individuals. This approach offers a unique opportunity to develop a non-invasive biosurveillance system for filoviruses that is both reproducible and compatible with field conditions, particularly in tropical African ecosystems (8).

This surveillance approach is particularly relevant in regions where Ebola virus has repeatedly emerged. The Democratic Republic of the Congo (DRC) has experienced repeated Ebola virus disease outbreaks since 1976, many of which have occurred in remote forested regions where human populations live in close proximity to wildlife (2). Equateur Province, including areas near the Mabali Reserve, is part of this historical hotspot, suggesting long-term viral circulation within local ecosystems (2). This regional context strengthens the rationale for investigating silent exposure to Ebola virus in wildlife species sharing these habitats.

Based on this premise, we hypothesize that free-ranging NHPs may serve as true ecological sentinels, capable of reflecting viral dynamics in the environment, even in the absence of apparent outbreaks. Using an innovative approach, fecal serology studied by an automated western blot system (JESS), we aimed to explore the seroprevalence against EBOV in wild NHPs from the Ebola fever endemic region, in Mabali Reserve, in Equateur Province (DRC).

## 2. Materials and Methods

### 2.1. Study site and sample collection

Between December 1 and 20, 2023, a total of 630 fecal samples from NHPs were collected in the Mabali Forest Reserve, located in Equateur Province (DRC) (Figure 1).

**Figure 1.**
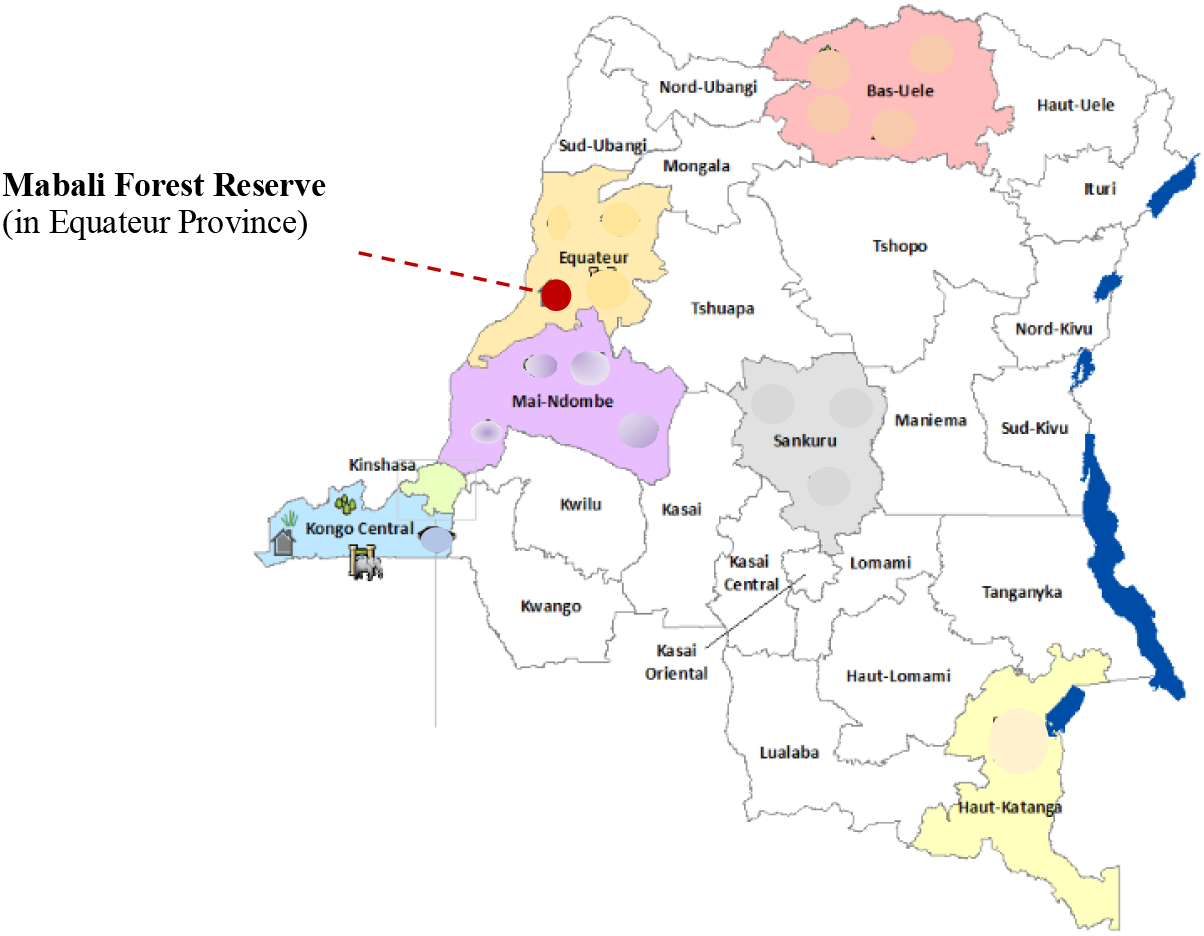
Geographic location of the Mabali Forest Reserve in Équateur Province, Democratic Republic of Congo.

Feces were collected directly from the ground in their natural habitat, without any direct contact with the animals. Animal origin of feces was evaluated macroscopically by experienced field guides. Each sample was individually placed in a hermetically sealed Ziploc-type bag and immediately stored on-site in a −80 °C freezer.

### 2.2. Transport and storage

At the end of fieldwork, samples were transported to the National Institute of Biomedical Research (INRB) in Kinshasa using a portable freezer maintaining a constant temperature of −80 °C. They were temporarily stored at the INRB before being shipped to the Institut Hospitalo-Universitaire Méditerranée Infection (IHU-MI) laboratory in Marseille (France). International transport was carried out using dry ice, ensuring preservation of the cold chain throughout shipment.

### 2.3. Ethical approval

The collection of NHP samples was conducted in accordance with ethical regulations and authorized by the Center for Research in Ecology and Forestry (CREF), Mabali, under the approval letter No. 0051/MIN.R.S.I.T./CREF/MAB/DG/01/MMIK/2023, issued on 17 November 2023. Authorisation of import.

### 2.4. Sample preprocessing

Upon arrival at the IHU-MI laboratory, each sample was divided into two 200 mg aliquots and one 2 g aliquot. Due to previous Ebola outbreaks in the study region, the first aliquot was subjected to an inactivation protocol for potential Ebola virus, following the method described by Haddock et al. (2016) (16).

### 2.5. Nucleic acid extraction and molecular detection of Ebola virus (First aliquot)

After inactivation, samples were homogenized and incubated for ≥ 4 h with proteinase K and NucliSENS easyMAG® lysis buffer (bioMérieux, Marcy-l’Étoile, France). Combined RNA/DNA extraction was performed using the NucleoMag Tissue kit (Macherey-Nagel, Düren, Germany) on the KingFisher Flex System (Thermo Scientific, Vantaa, Finland). Extracts were stored at −20 °C until PCR analysis. A qPCR assay targeting the nucleoprotein (NP) gene of Ebolavirus (Table 1) was performed to confirm the absence of viral material prior to further processing, using the CFX96 Real-Time PCR System (Bio-Rad, Hercules, CA, États-Unis). The total reaction volume (20 µL) contained: 5 µL of template RNA, 5 µL of Fast Virus 1-Step Master Mix (Thermo Fisher Scientific, Waltham, MA, États-Unis), 0.5 µL of each primer (10 pmol/µl), 0.4 µL of FAM-labeled probe (10 pmol/ µl) and 8.6 µL of DNase/RNase-free water. Thermal profile: 50 °C for 10 minutes for reverse transcription, 95 °C for 30 seconds for pre-denaturation, followed by 45 cycles of 95 °C for 5 seconds and 60 °C for 30 seconds.

**Table 1.**
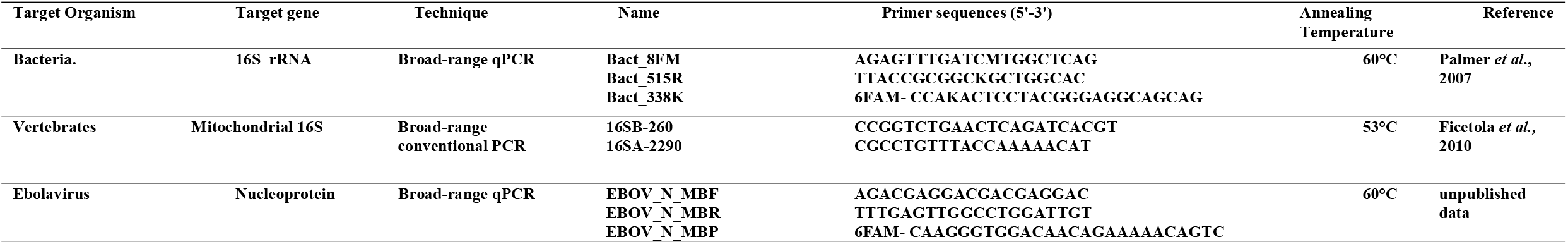
Oligonucleotide sequences of primers and probes used for real-time PCR and conventional PCR in this study.

### 2.6. Nucleic acid extraction (second aliquot)

The second non inactivated aliquot of each sample was also subjected to combined RNA and DNA extraction, following exactly the same protocol as for the first aliquot. The extracts were stored at –20 °C until molecular analysis.

### 2.7. Verification of bacterial DNA quality

Before implementing targeted molecular screening, a universal qPCR targeting the 16S rRNA gene (Table 1) was performed on all fecal samples (second aliquot) from NHPs to verify the presence of bacterial DNA and assess extraction quality. It was performed in a final volume of 20 µL containing 5 µL of template DNA, 10 µL of Roche Master Mix, 0.5 µL of each primer (20 µM), 0.5 µL of FAM-labeled probe (5 µM), and 3.5 µL of DNase/RNase-free distilled water. Reactions were run on a CFX96 Real-Time PCR thermocycler (Bio-Rad, Hercules, CA, États-Unis) under the following thermal program: 10 minutes at 95 °C, followed by 39 cycles of 10 seconds at 95 °C and 30 seconds at 60 °C. This step confirmed the biological origin of the samples collected in the field, excluded those with low bacterial load or degraded DNA, and facilitated the detection of potential PCR inhibitors. Extracted nucleic acids were diluted tenfold in sterile distilled water prior to being used for molecular identification of NHP species and screening for other microorganisms, with the goal of optimizing the performance of subsequent PCR reactions.

### 2.8. Molecular identification of primate species

Molecular identification of NHP species was performed using conventional PCR targeting the mitochondrial 16S DNA gene (Table 1). Reactions were carried out on a GeneAmp PCR Systems thermocycler (Applied Biosystems, Foster City, CA, États-Unis), in a final volume of 25 µL containing 5 µL of template DNA, 12.5 µL of Ampli Taq Gold Master Mix, 0.75 µL of each primer (20 µM), and 6 µL of DNase/RNase-free distilled water. The thermal program included an initial step at 95 °C for 15 minutes, followed by 35 cycles of 30 seconds at 95 °C, 30 seconds at 53 °C, 1 minute at 72 °C, and a final extension of 7 minutes at 72 °C. Amplification products were visualized by electrophoresis on 1.5% agarose gel. Samples showing a ∼600 bp band were considered positive and were purified using NucleoFast 96 PCR plates (Macherey–Nagel, Düren, Germany) before sequencing.

### 2.9. Serological analysis (Third aliquot)

The third aliquot was processed for serological assays following a modified protocol from Sereme et al. (2021) (17). For each sample, 2 g of feces were resuspended in 5 mL of PBS containing protease inhibitors (Pierce™ Protease Inhibitor Tablets, Thermo Fisher Scientific, Waltham, MA, USA), vortexed for 1 minute, and incubated at room temperature for 1 hour. Samples were then centrifuged at 2,500 rpm for 90 min at 4 °C, and the supernatant was filtered using a 0.22 µm membrane. The filtrate was lyophilized for 24 h at −80 °C and subsequently reconstituted in 500 µL of cold distilled water. The final protein concentration was determined using the Amicon® Ultra-4 centrifugal filter units (Millipore, Burlington, MA, USA).

### 2.10. Automated Western blot (JESS)

Concentrated proteins were analyzed using an automated capillary-based immunoassay system (Jess™, ProteinSimple, San Jose, CA, USA) to detect total anti-Ebolavirus immunoglobulins. Recombinant EBOV antigens (nucleoprotein (NP), glycoprotein (GP), viral protein 40 (VP40)) were obtained from Sino Biological (Eschborn, Germany). NP (Cat# 40443-V07E1) and VP40 (Cat# 40446-V07E) were expressed in *Escherichia coli*, whereas GP (Cat# 40442-V08B1) was expressed in baculovirus-infected insect cells. All proteins corresponded to Zaire ebolavirus (strain H.sapienswt/GIN/2014/Kissidougou-C15) recombinant constructs with polyhistidine tags. Antigens were prepared at a final concentration of 200 µg/mL in Reagent Diluent consisting of 0.1X Sample Buffer and a Master Mix prepared according to the manufacturer’s instructions (including Sample Buffer 10X and DTT; ProteinSimple, Bio-Techne, San Jose, CA, USA), and subsequently diluted 1:500 before use. A validated positive control (survivor serum) was included to define the interpretation threshold. Samples were considered positive when a discrete band was detected at the same migration level as the positive control. NP (∼22 kDa) was used as the initial screening antigen. NP-reactive samples were subsequently analyzed for reactivity to GP (∼69 kDa) for confirmatory purposes. Only double NP/GP reactivity was interpreted as indicative of the presence of anti-Ebola antibodies. Detection of VP40 (∼69 kDa) constituted an additional element supporting the observed serological profile.

## 3. Results

### 3.1. Screening for Ebola Virus and Quality of DNA Extracts

Molecular screening for Ebolavirus, targeting the nucleoprotein gene, was conducted on the extracts obtained from the first aliquot. The results confirmed that all samples (630/630; 100%) were negative, thereby allowing their processing under standard biosafety level 2 conditions (BSL-2).

To assess the quality of nucleic acid extraction from the second aliquot, a qPCR targeting the bacterial 16S rRNA gene was performed on 243 randomly selected samples. High Ct values (≥ 33) were observed in the crude extracts, indicating the presence of PCR inhibitors. A 1:10 dilution was applied, which successfully reduced Ct values to < 30. Consequently, this dilution was systematically applied to all subsequent molecular analyses.

### 3.2. Molecular Identification of NHP Species

PCR amplification of the mitochondrial 16S gene on all 630 diluted DNA extracts achieved a molecular identification success rate of 90.3% (569/630). The species *Cercopithecus ascanius* was predominant, representing 62.2% of identified samples. Detailed identity percentages and GenBank accession numbers are presented in Table 2.

**Table 2.**
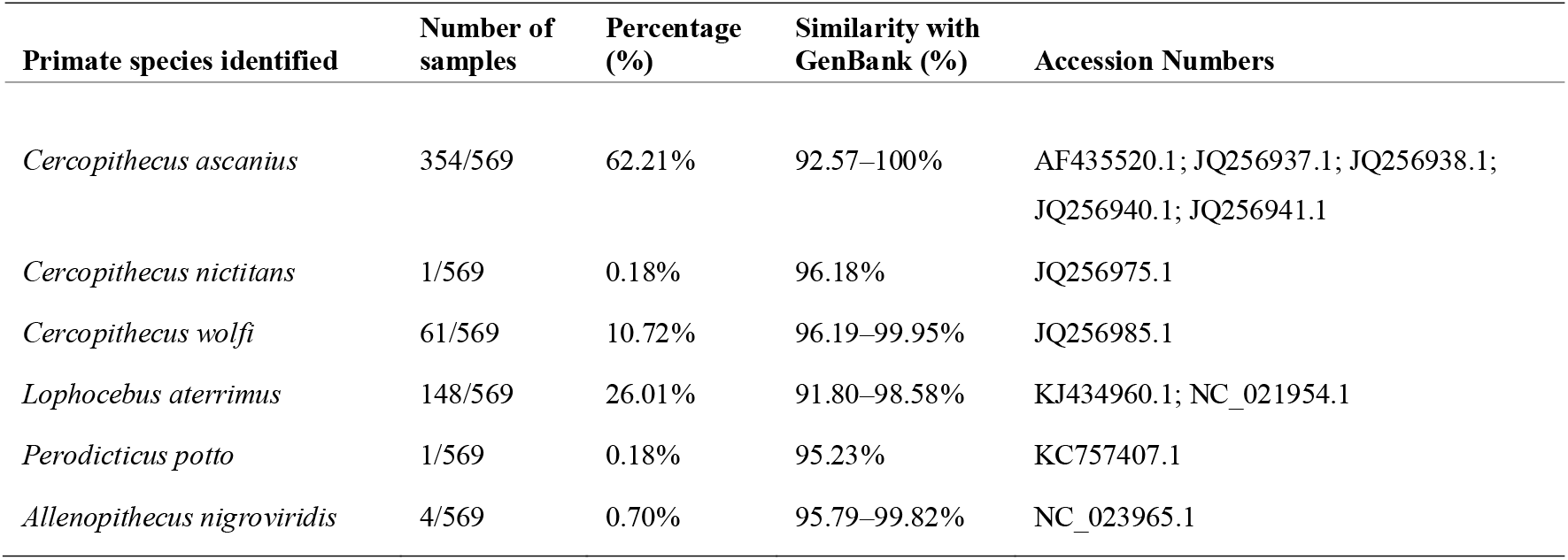
Primate species identified by fecal barcoding.

| Primate species identified | Number of samples | Percentage (%) | Similarity with GenBank (%) | Accession Numbers |
| --- | --- | --- | --- | --- |
| *Cercopithecus ascanius* | 354/569 | 62.21% | 92.57–100% | AF435520.1; JQ256937.1; JQ256938.1; JQ256940.1; JQ256941.1 |
| *Cercopithecus nictitans* | 1/569 | 0.18% | 96.18% | JQ256975.1 |
| *Cercopithecus wolfi* | 61/569 | 10.72% | 96.19–99.95% | JQ256985.1 |
| *Lophocebus aterrimus* | 148/569 | 26.01% | 91.80–98.58% | KJ434960.1; NC_021954.1 |
| *Perodicticus potto* | 1/569 | 0.18% | 95.23% | KC757407.1 |
| *Allenopithecus nigroviridis* | 4/569 | 0.70% | 95.79–99.82% | NC_023965.1 |

### 3.3. Serological Analyses

Serological analyses were performed using protein extracts from the third aliquot, following the protocol described in Section 2.9. A total of 569 samples from molecularly identified NHPs were analyzed using the automated western blot system (JESS) to detect total anti-Ebolavirus immunoglobulins (Ig). Four samples (4/569; 0.70%) were reactive to NP and GP (Figure 2) and additionally to VP40, consistent with seropositivity. These positive samples originated from two *C. ascanius* (2/354; 0.56%) and two *C. wolfi* (2/61; 3.27%). No seroreactivity was detected in *Lophocebus aterrimus* (0/148), *Allenopithecus nigroviridis* (0/4), *C. nictitans* (0/1), or *Perodicticus potto* (0/1) (Table 3).

**Table 3.** Detection rates of total anti–Ebola virus immunoglobulins in non-human primates.

| Antigen | NHP species |  |  |  |  |  | Total<br>N=569 |
| --- | --- | --- | --- | --- | --- | --- | --- |
|  | *Cercopithecus*<br>*ascanius*<br>N=354 | *Lophocebus*<br>*aterrimus*<br>N=148 | *Cercopithecus*<br>*wolffi*<br>N=61 | *Allenopithecus*<br>*nigroviridis*<br>N=4 | *Cercopithecus*<br>*nictitans*<br>N=1 | *Perodicticus*<br>*potto*<br>N=1 |  |
| NP EBOV Ag | 2 (0,56 %) | 0 (0%) | 2 (3,27 %) | 0 (0 %) | 0 (0 %) | 0 (0 %) | 4 (0,70 %) |
| VP40 EBOV Ag | 2 |  | 2 |  |  |  |  |
| GP EBOV Ag | 2 |  | 2 |  |  |  |  |

**Figure 2.**
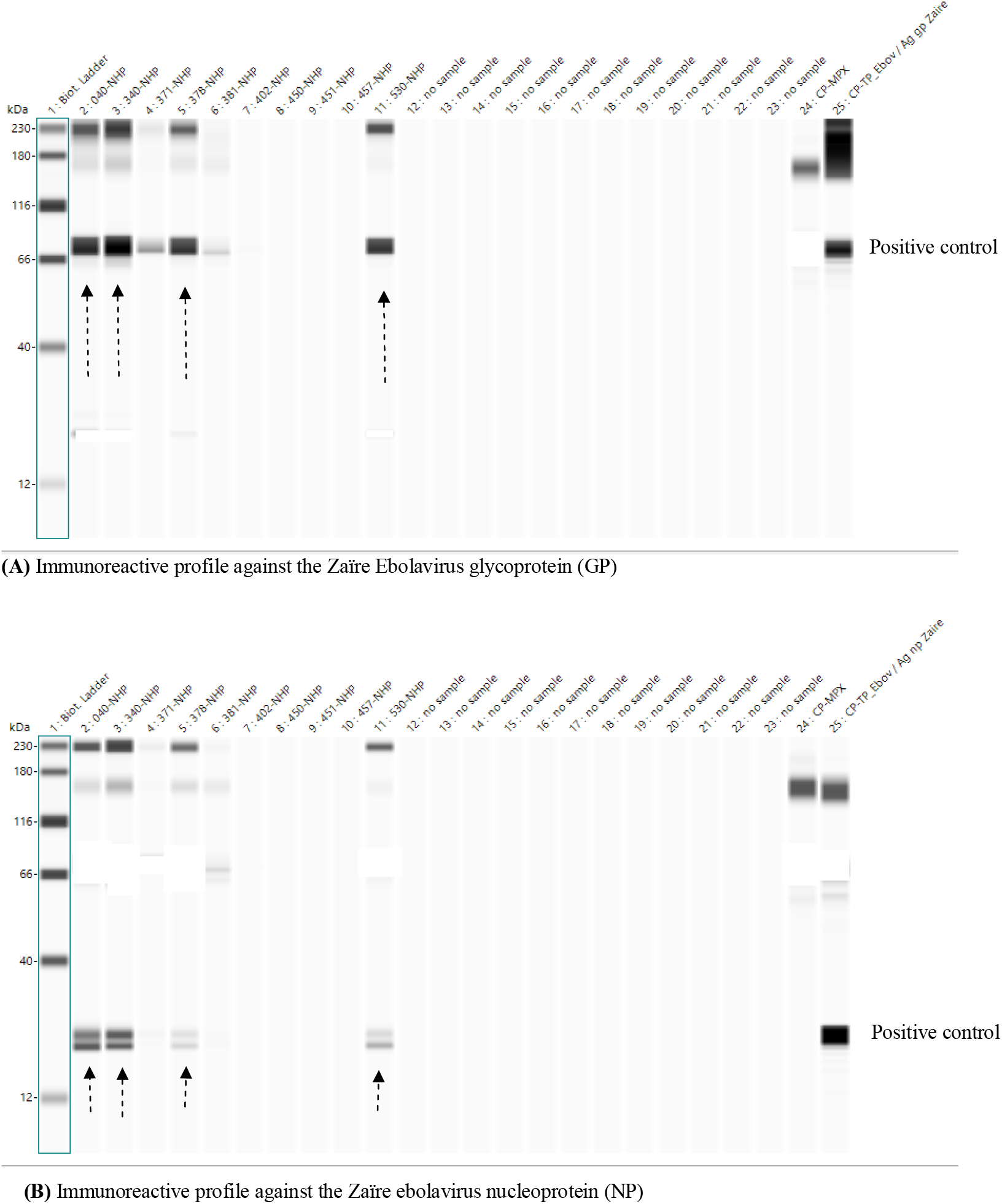
Serological immunoreactivity against Zaïre ebolavirus antigens assessed by automated capillary Western blot using the Jess system (ProteinSimple).

## 4. Discussion

None of the 630 fecal samples tested showed a PCR-positive signal targeting the nucleoprotein (NP) gene of Ebolavirus. This result is consistent with the absence of an officially reported outbreak in the Équateur Province at the time of sampling, although the region has historically experienced several epidemic events (18). However, the absence of molecular detection cannot be interpreted as proof of viral absence. Indeed, several studies have shown that filoviruses may circulate according to intermittent (7) or cryptic dynamics (19), thereby eluding classical PCR-based detection methods. Therefore, PCR alone is insufficient to exclude the possibility of a low-level viral circulation within the investigated ecosystem.

This interpretation is further supported by field studies that reported negative PCR results in African nonhuman primates, despite serological evidence of prior exposure to Ebolavirus. This was particularly observed in fecal samples from wild great apes in Gabon (20)(8), where the absence of molecular detection was accompanied by specific immune responses, suggesting that past infections may fall below the detection threshold of current PCR assays. Thus, molecular negativity should not be equated with viral exclusion from the ecosystem.

Conversely, PCR-positive results reported in the literature largely originate from actively infected individuals, carcasses collected during or immediately after an outbreak (21)(22), or experimental infection models in macaques, in which viral loads are considerably higher (23). Under these specific conditions, molecular detection corresponds to clinical data. This contrasts strongly with non-invasive surveillance strategies conducted during inter-epidemic periods, particularly those using fecal samples, for which serological approaches frequently prove to be more sensitive than PCR (20)(8).

These findings highlight the necessity of complementing molecular approaches with non-invasive serological methods capable of identifying immune signatures of past exposure, even in the absence of active infection. In this context, fecal serology represents a methodologically relevant alternative (17), particularly well-suited to free-ranging African nonhuman primates.

Although the seroprevalence observed in our study was low (0.70%, i.e., 4/569), the detection of a ∼22 kDa immunoreactive signal, compatible with the Ebolavirus nucleoprotein, represents a noteworthy eco-epidemiological observation. This signal may correspond to a truncated form or a degradation product of NP, as previously suggested by experimental studies (24). The occurrence of this band exclusively in *C. ascanius* (2/354; 0.56%) and *C. wolfi* (2/61; 3.27%) raises the possibility of discrete environmental exposures or circulation of filovirus-related agents, potentially below the thresholds of molecular detection.

Similar seroprofiles, of low intensity but with biological significance, have been reported in other wild African primates. Leroy et al. (2004) (21) documented seropositivity in wild chimpanzees (up to 12.9%), several *Cercopithecus* species (6.7%), and Mandrillus (2.8%) using ELISA followed by confirmatory Western blot targeting multiple Ebola virus antigens including nucleoprotein (NP) and glycoprotein (GP). Likewise, Reed et al. (2014) (20) demonstrated the feasibility of detecting anti-EBOV antibodies in nonhuman primate feces using indirect ELISA targeting EBOV NP and GP, and Mombo et al. (2020) (8) used similar serological assays on fecal extracts, with antibodies directed against EBOV NP and GP, to detect exposure in great apes. These findings indicate that Ebolavirus infection is not necessarily fatal in wild primates and that immune signatures may persist long after epidemic events, even when molecular detection is negative.

The relatively low seroprevalence observed in our study is likely attributable to several complementary factors. First, we adopted a conservative, fully non-invasive strategy based on automated western blot analysis (JESS), targeting total immunoglobulins and applying a strict positivity threshold (≥ 3× the negative control). Importantly, automated capillary immunoassays require only minimal amounts of protein extract, a key advantage when working with wildlife fecal samples where sample material is limited and often degraded. In addition, automated capillary-based systems such as Jess Simple Western™ provide high analytical specificity by generating discrete molecular weight peaks rather than broad, non-specific signal intensities, thereby reducing cross-reactivity and false-positive interpretations compared with traditional Western blots (25). This analytical framework favors specificity over sensitivity, limiting false positives and enhancing reproducibility (17). Consistent with this specificity-oriented approach, the interpretation criteria relied on a conservative sequential algorithm: an initial screening step based on nucleoprotein reactivity, followed by confirmatory testing against glycoprotein for NP-reactive samples. Only samples exhibiting double NP/GP reactivity were classified as seropositive, with VP40 detection considered supportive evidence. While this stringent algorithm enhances analytical specificity and reduces the risk of cross-reactivity or false-positive interpretation, it may have limited sensitivity and potentially underestimated low-titer or partially degraded antibody responses in fecal samples. It cannot be excluded that some samples reactive exclusively to GP may not have been detected as a result of this sequential screening strategy. Second, the four positive samples originated exclusively from *C. ascanius* and *C. wolfi*, opportunistic frugivorous species inhabiting forest edges and forest–open area ecotones (26)(27). Unlike gorillas or chimpanzees, which may engage in scavenging or carcass manipulation, these species are primarily frugivorous, which may reduce their exposure to infectious tissues and direct viral transmission, potentially explaining their comparatively low seroprevalence (13).

To our knowledge, no previous study has reported Ebolavirus seroprevalence estimates in *C. ascanius* or *C. wolfi*. Although sporadic seropositive cases have been documented in other *Cercopithecus* species (21)(28), our findings would therefore represent the first immunological evidence of potential exposure in these two taxa. This observation suggests a possible eco-epidemiological role for the genus Cercopithecus in filovirus dynamics, particularly in ecotones and forest–anthropized interfaces. Given their high abundance in fragmented forests, opportunistic frugivory, and tolerance to habitat disturbance, these species may function as ecological sentinels or bridge hosts, even in the absence of apparent outbreaks. Detecting a low but measurable seroprevalence in *C. ascanius* and *C. wolfi* may thus represent a valuable epidemiological signal of low-level viral circulation, supporting the integration of these species into One Health surveillance frameworks.

Our sampling was conducted during an inter-epidemic period in a relatively preserved and low-anthropized forest environment. Yet several studies have shown that forest fragmentation, anthropogenic pressure, and environmental disturbances significantly increase the risk of zoonotic spillover (29)(30)(31). In addition, Olival et al. (2017) (32) demonstrated that integrating host traits, viral characteristics, and anthropogenic variables can predict hotspots of zoonotic potential, reinforcing the value of targeted surveillance in primates inhabiting ecological interfaces. Therefore, our low seroprevalence may represent not a lack of viral circulation, but rather an ecological baseline characteristic of a “silent” period, an informative reference for the implementation of longitudinal surveillance efforts. However, the mortality rate associated with filovirus infection in *Cercopithecus* species remains largely unknown, and it cannot be excluded that higher mortality could also contribute to the low proportion of seropositive individuals detected.

Finally, minimizing the risk of serological cross-reactivity was a major strength of our methodological approach. The systematic exclusion of non-specific bands, targeted detection of NP, GP and VP40 antigens, and the use of automated JESS technology helped prevent erroneous interpretations due to cross-reactivity (24). Collectively, these methodological considerations indicate that the low seroprevalence observed in our study likely reflects improved diagnostic specificity and provides a robust foundation for eco-epidemiological monitoring of filoviruses in African nonhuman primates.

## 5. Conclusion

To the best of our knowledge, this study provides the first evidence of seropositivity compatible with exposure to Ebolavirus in *C. ascanius* and *C. wolfi*, two opportunistic species inhabiting ecological interfaces between forest and anthropized areas. Although overall prevalence was low (0.70%), detection of a specific immunoreactive signal represents a significant indication of discrete viral circulation within the surveyed ecosystem. These findings support the hypothesis of a potentially intermittent, cryptic or low-level endemic viral dynamic that may escape PCR-based detection during inter-epidemic periods.

The non-invasive methodological framework applied here, particularly fecal serology using the automated JESS western blot system, demonstrated relevance for investigating highly zoonotic pathogens in ecological surveillance contexts. By reducing biases associated with capture, stress, or serum sampling, this approach offers a perspective on past exposures, allowing detection of viral circulation beyond clinically observable outbreaks.

This first serological characterization within the Cercopithecus genus opens perspectives for future research, including longitudinal monitoring to assess spatial, temporal, and seasonal variations in seroprofiles; integration of ecological indicators (forest fragmentation, land use, human density) into predictive models of zoonotic risk; combining non-invasive serology with metagenomics and behavioral data to better identify potential exposure routes.

Within a One Health framework, NHPs, especially generalist species such as *C. ascanius* and *C. wolfi*, appear to be ecological sentinels for monitoring filoviruses in tropical forest ecosystems. The low seroprevalence observed should therefore not be interpreted as lack of risk, but rather as an eco-epidemiological baseline during an inter-epidemic period. In the context of increasing environmental disturbances across Central Africa, implementation of non-invasive surveillance strategies, including fecal serology, is feasible and necessary to anticipate future emergence events.

## Acknowledgements

The authors gratefully acknowledge Simon-Pierre Ndimbo and the team of the Centre de Recherche en Écologie et Foresterie (CREF) of Mabali for their field support. We also sincerely thank Ms. Marielle Bedotto for their essential contributions to the successful conduct of this study.

## Notes

### Competing Interest Statement

The authors have declared no competing interest.

